# Field-based evidence of sperm quality impairment associated with conventional farming in two passerine birds

**DOI:** 10.1101/2025.09.17.676817

**Authors:** Ségolène Humann-Guilleminot, Audrey Bailly, Karine Monceau, Mélissa Desbonnes, Vincent Bretagnolle, Jérôme Moreau

## Abstract

The detrimental effects of conventional farming on bird biodiversity are increasingly documented. Despite this, the specific impacts of both organic and conventional farming practices on bird coloration and sperm quality in natural settings remain unexplored. This study aimed to determine whether these farming practices differentially affect body mass, coloration intensity, and sperm quality in passerine birds inhabiting agricultural landscapes. We captured seven passerine species in hedgerows adjacent to fields farmed organically or conventionally, within a 250-meter radius representative of their breeding home ranges. Body mass was measured across all species, and an index of coloration intensity, based on carotenoid (yellow/orange) and melanin (black) pigments, was assessed in four species. Additionally, we evaluated three sperm quality parameters (sperm density, percentage of abnormal sperm, and intra-individual variance of sperm morphology) using fresh sperm samples collected from four species in the field. We hypothesized that birds living near conventional fields would exhibit lower body mass due to reduced food availability. Additionally, we predicted that the more favourable conditions associated with organic farming, such as greater food abundance and lower exposure to pollutants, would benefit birds, leading to enhanced coloration and improved sperm quality. Our results did not reveal any difference in body mass or coloration. The absence of observable effects might be due to several factors: methodological limitations, cross-contamination between habitats or insufficient exposure to farming practices that may hide any potential difference between the two habitats, or the intrinsic adaptive strategies of the species. However, subgroup analysis of three and four species revealed a decrease in sperm density and a higher proportion of abnormal spermatozoa in the Common Whitethroat and Common Nightingale living in conventional farming, respectively. Although our sample size was limited, we believe these findings highlight the potential negative effects of conventional farming on birds.

## 1. Introduction

Agricultural lands cover nearly 40% of the world’s terrestrial surface and its intensification has been associated with a significant threat to global biodiversity worldwide (BirdLife International, 2022; Emmerson et al., 2016; Flohre et al., 2011; Karp et al., 2012; Stoate et al., 2009). Agricultural intensification notably involves the replacement of natural ecosystems into monocultures, marked by intensive pesticide use, over-fertilization, and the alteration of natural landscapes, which lead to soil erosion, pollution, and a decrease in habitat quality and biodiversity (Burns et al., 2016; Christel et al., 2021; Emmerson et al., 2016; Flohre et al., 2011; Sánchez-Bayo and Wyckhuys, 2019). The rate of biodiversity loss has dramatically increased over the last century, particularly impacting avian species, that are key indicators of ecological health. This decline has been linked to agricultural intensification (Rigal et al., 2023; Stanton et al., 2018), and may be driven by various changes in agricultural practices impacting bird populations through multiple pathways. First, the replacement of diverse crops interspersed with grasslands by intensively cultivated crops not only diminishes available food sources for many species but also reduces nesting opportunities, subsequently increasing competition among and within species (Assandri et al., 2017; Beecher et al., 2002; Kragten et al., 2011). Second, the application of chemicals such as pesticides can directly poison birds, resulting in either immediate lethal effects or chronic sublethal impacts on various physiological parameters (Gibbons et al., 2015; Humann-Guilleminot et al., 2024b, 2023; Mineau et al., 2001; Richard et al., 2021).

To mitigate the impact of agriculture on biodiversity, several European measures have been implemented under the Common Agricultural Policy (CAP; European Commission, 2022). These include ecological focus areas which help maintain attractive landscapes and are extensively or low-intensively managed (e.g., hedgerows, ponds, extensive meadows), and in-production schemes that support environmentally friendly crop production such as organic farming (Batáry et al., 2015; European Commission, 2010). Under European Union regulations, organic farming very strictly restricts the use of synthetic products on crops, prompting farmers to adopt agro-ecological methods that promote crop diversity and environmentally friendly cultivation practices (European Commission (EU) 2018/848). Several studies have shown that organic farming increases the abundance and diversity of weeds and invertebrates, therefore increasing the availability of food for birds, but also enhances overall habitat quality and minimizes the risk of pesticide poisoning (Beecher et al., 2002; Chamberlain et al., 1999; Gayer et al., 2021; Gong et al., 2022; Hole et al., 2005). Notably, a study reported more vigorous behaviours from birds captured in organic farming compared to birds captured in conventional farming (Moreau et al., 2022a). On the contrary, agricultural intensification has been shown to adversely affect corn bunting breeding success by decreasing the availability of invertebrate food sources due to pesticide use and inadequate foraging habitats (Brickle et al., 2000). Another study found that House sparrows living in conventional farms were lighter than birds living in organic and integrated-production farms (Humann-Guilleminot et al., 2024a). Furthermore, many chemicals commonly used in conventional agriculture, especially pesticides, can impair bird sperm quality through direct damage to sperm cells by inducing oxidative stress or acting as endocrine disruptors. These substances can lead to testicular anomalies or reduced spermatozoa production, potentially diminishing reproductive success (Humann-Guilleminot et al., 2019b; Khalil et al., 2017; Mohanty et al., 2017; Moreau et al., 2022b). Additionally, hormonal changes associated with plumage depigmentation were observed in male Red Avadavats exposed to imidacloprid and mancozeb (Pandey et al., 2017).

Ornamental traits evolve with sexual selection, revealing the quality and condition of their bearer (Cotton et al., 2004) and their plasticity makes their expression highly sensitive to environmental stressful factors (Buchanan, 2000; Hill, 1995; Lifshitz and St Clair, 2016). The two primary types of pigments involved in the coloration of ornamental traits in vertebrates are melanin and carotenoids. Melanin are the most prevalent pigments found in vertebrates, with pheomelanin producing yellow-brown features, and eumelanin producing grey-black features (McGraw, 2005). Melanin-based ornaments are mainly controlled by genes (synthesized from amino acid precursors, (Badyaev and Hill, 2000; McGraw, 2006a), but they can also be affected by environmental conditions such as breeding conditions, parasite infestations and food quality (Fargallo et al., 2007; Guindre-Parker and Love, 2014; McGraw, 2008). The other type of pigment, carotenoids, produce yellow, orange, and red traits. They cannot be synthesized in vertebrates (Schiedt, 1989), but they are acquired through the diet. They can therefore supply information on foraging, carotenoid absorption and efficiency of resource allocation (McGraw, 2006b). They are involved in some physiological functions, preserving cells and tissues from oxidative damage, and strengthening the immune system (Hartley and Kennedy, 2004; Lozano, 1994; Pérez-Rodríguez, 2009; Schantz et al., 1999). Both melanin and carotenoids are relevant for assessing the negative effects of pesticides on physiological parameters in wildlife. For example, exposure of Red-legged Partridges to an herbicide led simultaneously to an increase of eumelanic plumage (larger black traits), a decrease in pheomelanin expression (smaller brown traits) and also to a decrease in carotenoid-based ornaments (paler red beak and eye rings; Alonso-Alvarez and Galván, 2011).

In this study, we aimed to explore the effects of conventional and organic farming on birds’ body mass, birds’ coloration intensity and sperm quality parameters in birds captured in hedgerows within landscapes surrounded by either organically or conventionally farmed fields. For this purpose, we examined carotenoid-based (yellow/orange) and melanin-based (black) coloration across four bird species from which we derived an index of colour intensity. In addition, we selected sperm density, percentage of abnormal sperm and intra-individual variation of sperm morphology as proxies for sperm quality in eight passerines species. The choice of these sperm quality parameters relies on several variables that, individually or in combination, have been demonstrated to effectively predict the fertilizing potential of sperm across various species (Simmons and Fitzpatrick, 2012; Snook, 2005). We opted to explore intra-individual variation in sperm morphology, i.e., variation in total sperm length within ejaculates of each individual, rather than absolute measurements of sperm components (e.g., head, midpiece) due to the diversity in reproductive strategies among the nine passerine species studied, which range from low to high levels of sperm competition potentially affecting sperm quality and size variation. First, we hypothesized that farming practices may influence food abundance and therefore that birds living within conventional fields would be lighter. Second, we predicted that organic farming could enhance both bird coloration and sperm quality, potentially due to a lower exposure to pollutants or increases in food abundance and quality that are associated with improved overall body condition, which in turn is linked to sperm quality and coloration (Fernández-Eslava et al., 2022; Helfenstein et al., 2010; McDiarmid et al., 2022; Schantz et al., 1999).

## 2. Material and methods

### 2.1. Study design

The study was conducted in the ‘Zone Atelier Plaine & Val de Sèvre’ (ZA-PVS), a 435 km^2^ LonglTerm SociallEcological Research (LTSER) site in central-western France (46°23′N, 0°41′W; see Bretagnolle et al., 2018 for details on our site). Globally, the majority of the area is typical intensive farmland, made up of open landscapes with mainly winter cereals and other arable crops (maize, sunflower, oilseed rape, pea), temporary grasslands (such as alfalfa and clover) and permanent meadows. The site has a rather high diversity of types of farming system, with ca. 15 farms managed using conservation agricultural practices, ca. 80 farms using organic methods, and 300 using conventional methods, of which about 50% are mixed dairy/cereal farms. Agricultural land use within the site has been monitored annually at the field scale since 1994 and mapped on vectorlbased shapefiles (Bretagnolle et al., 2018). Every organically farmed field has been identified (see Wintermantel et al., 2019), providing a very detailed description of farming practices regarding pesticide use at field and landscape scales.

We used this detailed knowledge to select 10 hedgerows surrounded by organically farmed landscapes and 10 hedgerows surrounded by conventionally farmed landscapes, each with a 250-m radius buffer zone (the same 20 hedgerows that those described in Moreau et al., 2022a). Physical characteristics of hedgerows (such as total length, height, top width) are similar between conventional and organic farming (see Moreau et al., 2022a for details). The percentage of organic farming was calculated within this 250-m radius buffer. We chose this buffer-zone size to include the breeding home-range size for different passerine species (see below). For hedgerows defined as ‘organic hedges’, 73% to 98% of the buffer area was organically farmed, while ‘conventional hedges’ had almost no organic farming within the buffer area (see Table 1 in Moreau et al., 2022a for details).

**Table 1.**
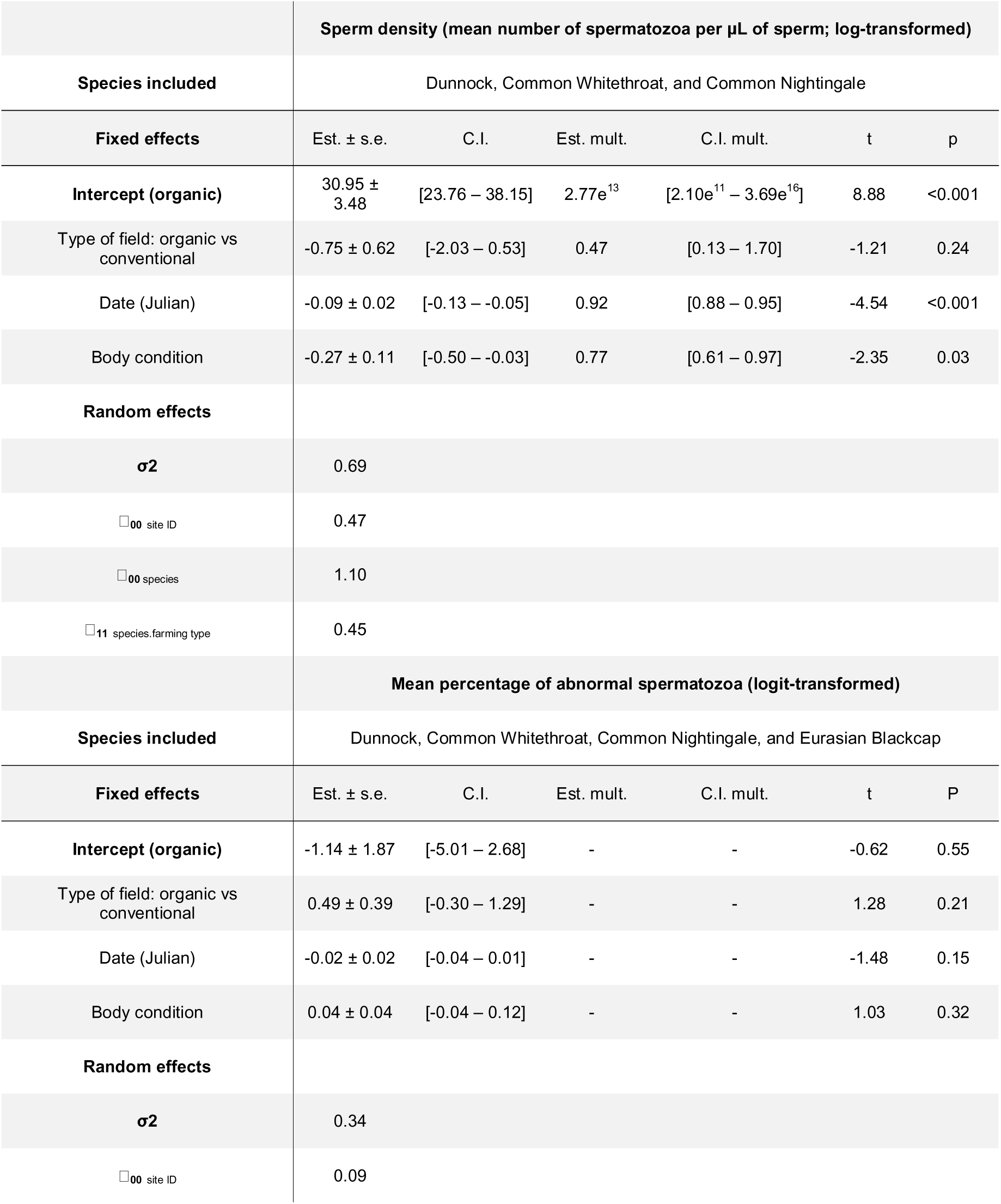
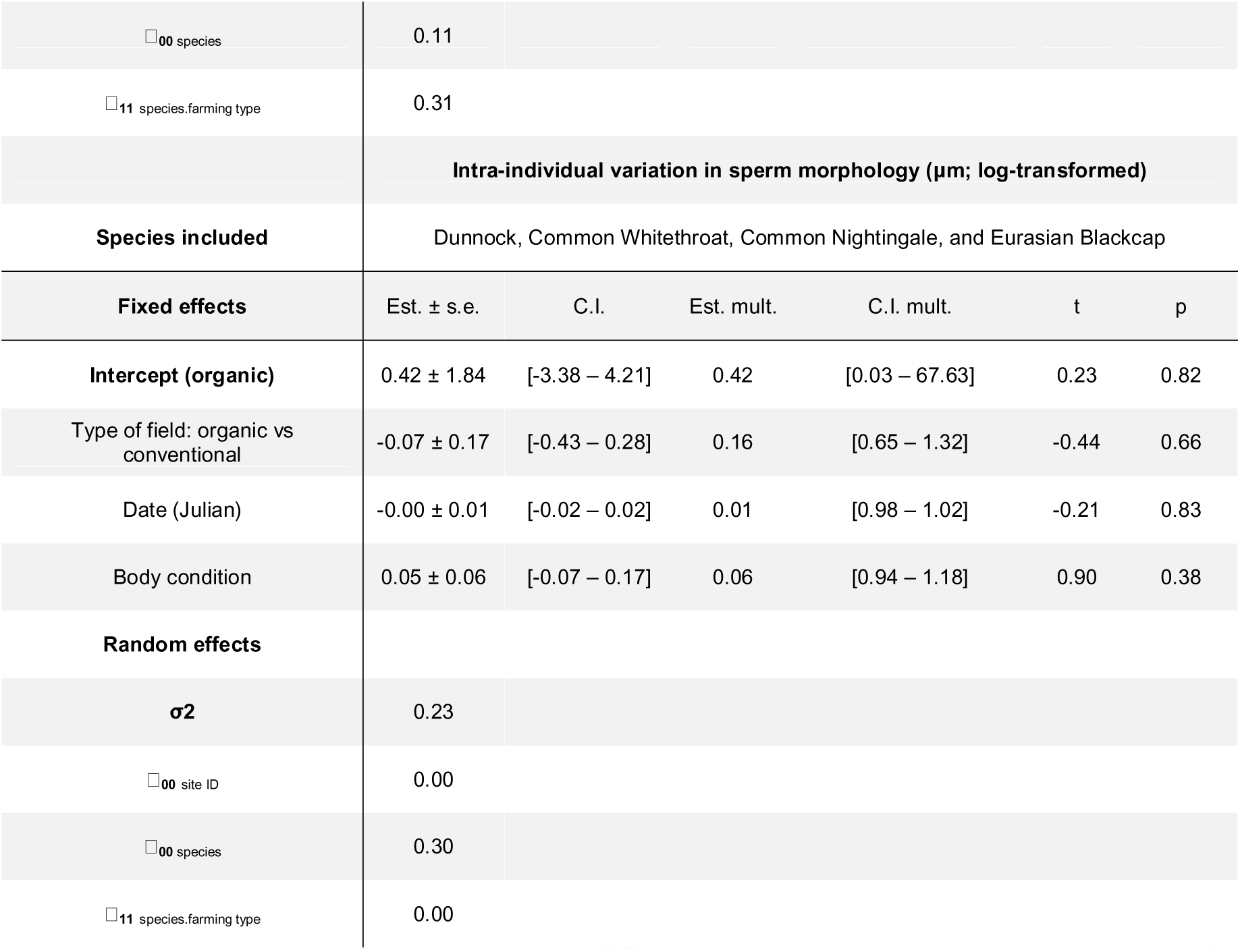
Summary of the linear models examining the impact of farming practices on sperm density, mean percentage of abnormal spermatozoa and intra-individual variation in sperm morphology in three to four species of birds fitted using a restricted maximum likelihood function. The table shows model estimates ± standard error (Est. ± s.e.) and associated 95% confidence intervals (C.I.); for log-transformed variables, the table also shows multiplicative estimates for untransformed variable (Est. mult.) and associated multiplicative confidence intervals (C.I. mult.); t-values and p-values. The multiplicative estimate of the intercept corresponds to the geometric mean of the variable.

The breeding territories of selected species vary significantly, spanning 0.2 to 8 hectares, equivalent to a radial expanse of 25 to 160 meters (Halupka et al., 2002; Naef-Daenzer, 1994; Naguib et al., 2001; Stevens et al., 2002; Stoate and Szczur, 2001). Notably, these breeding home ranges are more contained than the size of our study plots, which extend to a 250-meter radius, ensuring our analysis was conducted at an appropriate scale.

### 2.2. Bird capture

Adult male birds were captured using mist nets throughout the breeding season, from March 30, 2021, to July 19, 2021. Each site typically underwent three capture sessions, except for three conventional sites, where only two sessions were conducted due to bad weather conditions (rain and wind). Each site was visited one month apart, following the same order as during the first visit. Only one site was visited per day. For each capture, ten mist nets (each 12 meters long) were set up along hedges. The nets were placed early in the morning before sunrise until 11:30 am and in the evening until nightfall. They were checked every 5 to 10 minutes to ensure that any captured birds were promptly released, minimizing their time in the nets. In total, 28 different species were captured; however, the focus of this study was on seven targeted species: Cirl Bunting, Common Nightingale, Common Blackbird, Common Whitethroat, Eurasian Blackcap, Dunnock and the Great Tit. These species were selected based on the number of individuals caught for which there were sufficient amount of data for the assessment in birds’ body mass and coloration in the two types of sites. At capture, males were ringed and weighed using a spring balance (Pesola® 500 g, accuracy: ± 5 g). The left and right tarsus lengths were each measured twice with a digital caliper (precision: ± 0.1 mm). After all measurements done (see below), birds were released at the capture site immediately after measurement. Few birds were recaptured across the season for which only body mass was measured again. However, the small number of recaptures unevenly distributed across the two farms (N = 11) prevented us from assessing intra-individual changes over the breeding season. Only data from birds first capture was recorded in the database.

### 2.3. Ornamental colorations

Our study examined carotenoid-based (yellow/orange) and melanin-based (black) coloration across four bird species: the Great Tit, Cirl Bunting, Common Blackbird, and Eurasian Blackcap. We characterised colour on different parts of each of the four species. These parts were chosen because they are known to play an important role in intra- and intersexual communication and/or reflect the physiological quality of the individual in that species. For example, in Great Tits, individuals experimentally exposed to oxidative stress have less colourful carotenoid-based breast plumage (Helfenstein et al., 2010). We therefore assessed the yellow face lines of the Cirl Bunting, the black cap of the Eurasian Blackcap, the yellow breast plumage of the Great Tit and the orange bill of the Common Blackbird. Yellow colouration (bill or plumage) is carotenoid-based, and black plumage colouration is melanin-based.

Each bird was photographed twice within a black box using a ring light for consistent lighting, along with a standard white reference chip (Kodak Colour Separation Guide and Gray scale, Q13/Q14) for accurate colour calibration. The images were processed using PhotoFiltre software, where RGB values were measured from three random squares within the colour patterns and averaged across the two images per bird. These RGB values were then converted to HSB values using the *rgb2hsv* function from *grDevice* package (Fisher, 1999). Variation in light exposure assessed from the white reference chip (Fitze and Richner, 2002; Tschirren et al., 2003), was corrected to ensure accurate colour assessment. Principal Component Analysis (PCA) was then performed with the *FactoMineR* package (Husson et al., 2020) to generate a colour intensity index from the first principal axis, which reflected 67.20% of the variance and an eigenvalue greater than one. The first PCA axis was negatively correlated with the colour parameters H (Spearman’s rank correlation test: r = -0.83) and positively correlated with saturation and brightness (saturation r = 0.76, brightness r = 0.86). The mean of the first PCA axis values, calculated from two images per bird, provided an index of colour intensity, with higher scores indicating more intense coloration.

### 2.4. Sperm analyses

#### 2.4.1. Sperm collection

We attempted to collect ejaculates from all captured bird species without prior selection. However, sperm collection proved particularly challenging for some species, and we were able to retrieve ejaculates from four bird species: Eurasian Blackcap, Dunnock, Common Whitethroat and Common Nightingale. Sperm was collected by massaging the birds’ cloaca (Wolfson, 1952) until we reached a minimum sample volume of 2 μL. For some birds, continuous stimulation was needed to reach this amount, but the whole sperm sample was collected in the same 5 μL capillary tube (intraMark 5μL, BLAUBRAND®) and within 5 minutes of cloacal massage.

Immediately after sperm collection, we took a photograph of the capillary containing the ejaculate placed on a millimetre paper. Ejaculate size was then assessed using the ImageJ software as the length of the capillary section containing sperm times the squared radius of the capillary times π. Upon sperm collection, we also pipetted 0.20 μL of the ejaculate into 10 to 40 μL of ice-cold buffer (10% DMSO:90% new-born bovine serum), depending on the species and the density of the ejaculate, and the mixture was stored at -80°C until analysis of sperm density. In addition, a small droplet from the ejaculate was smeared with 10% formalin on a glass slide, dried, and stored at room temperature in the dark for further analyses on sperm morphology.

#### 2.4.2. Sperm quality parameters

We assessed sperm density, percentage of abnormal sperm and sperm morphology following the protocol described in (Humann-Guilleminot et al., 2019b).

##### 2.4.2.1. Sperm density

Sperm counts were conducted using Neubauer chambers, with a depth of 0.100 mm and a grid size of 0.0025 mm2. For the counting process, 1 μL of the sperm sample suspended in DMSO/serum was blended with 39 μL of PBS. In cases of high sperm concentration, an additional dilution was achieved by mixing another μL of the sample with 79 μL of PBS. Ten μL of the mixture was then loaded to the chamber slide. Spermatozoa were counted under a microscope set to 400x magnification with a phase-contrast 2 annular ring setting. The counting was focused on the nine primary squares of the chamber outlined by triple lines. We specifically counted sperm in 32 squares across two regions, each bounded by triple lines, including cells along the top and left borders while excluding those touching the right and bottom edges. Sperm density was then determined by dividing the total cell count by the chamber section volume of 0.004 μL and adjusted for the dilution level. We calculated the average sperm density from two separate measurements per individual. All chambers were cleaned with 70% ethanol and distilled water between each use. We obtained an intra-class correlation coefficient of 0.95 for the sperm counts and a coefficient of variation of 0.28, and we use the mean of two counts in the analyses.

##### 2.4.2.2. Sperm morphology

From each slide with the mixed sperm-formalin, we took photos of ten intact sperm cells using with a camera mounted on a microscope (Zeiss®, Primo star), at 200x or 400x magnification depending on species. Seven to 10 sperm cells per ejaculate (mean ± SD: 9.78 ± 0.57) were measured for head, midpiece, flagellum and total length using ImageJ software (v 1.53e). For further analyses, intra-male variation in sperm length was assessed by calculating the standard deviation of the mean total sperm length.

##### 2.4.2.3. Percentage of abnormal sperm

Sperm smears were also used to assess the percentage of abnormal sperm based on 50 spermatozoa per slide randomly selected. Spermatozoa were classified as morphologically normal or with one or more abnormalities (i.e., abnormal head – no head, S-shaped head, bended head, no acrosome, burst head, with abnormal midpiece – no midpiece, broken midpiece, abnormal flagellum – no flagellum, broken flagellum, folded flagellum, flagellum with 90° angle, coiled flagellum, double flagellum, split flagellum; (Humann-Guilleminot et al., 2018). We obtained an intra-class correlation coefficient of 0.81 for the two measurements and a coefficient of variation of 0.30, and we use the mean of two counts in the analyses.

### 2.5. Statistical analyses

All statistical analyses were performed using the R v. 4.2.2 software (R Core Team, 2022). We first investigated the effect of the two farming practices on birds’ body mass by running a linear mixed effect model (LMM) including the log-transformed mass of birds as the dependent variable and farming practices (organic vs conventional), bird species (seven species, categorical) and its interaction with farming practices, date (continuous numeric values in Julian date), tarsus length (mean of four measurements), and session (morning and evening, categorical) as explanatory fixed factors. Considering the correlation between birds’ body mass and tarsus length and to limit the number of variables in each model, we decided to use the scaled mass index according to (Peig and Green, 2009), as a proxy for birds’ body condition, instead of body mass and tarsus length separately. We then ran a LMM including the variation in coloration intensity (untransformed) as the dependent variable, farming practices, bird species and its interaction with farming practices, date, and scaled mass index as explanatory fixed factors. We conducted an analysis of variance using the *Anova* function from the *car* package, specifying a Wald chi-square test and a Type III sum of squares ANOVA. Post-hoc comparisons between farming practices within each species were performed using the *emmeans* function from the *emmeans* package (Lenth, 2023), with Tukey’s HSD adjustment to control for multiple comparisons.

We then ran three LMMs including either log-transformed sperm density, logit-transformed percentage of abnormal sperm or log-transformed intra-individual variation in sperm morphology as dependent variables and farming practices (organic vs conventional), date (continuous numeric values in Julian date), and scaled mass index as explanatory fixed factors. All three models included the site identity as random intercept to account for the non-independence of the samples taken in the same site. We did not include the session (morning or evening) as explanatory fixed factor in the models because sperm quality unlikely varies over the course of a day. Due to the low number of individuals caught within each species in the study of farming practices’ effects on sperm quality parameters, a full model fitting an interaction between species and farming practices would render insufficient degrees of freedom. Instead, we modelled species ID as a random effect with both random intercepts and random slopes for farming practices, thereby accounting for inherent differences among species and allowing each species to respond differently to farming practices. It allows for each species to have a different baseline response (random intercept) and a different reaction to the change of farming practices (random slopes). We acknowledge that combining all species into a single model could introduce excessive variance and potentially hide differences in sperm quality between habitats. To address this, we conducted subgroup analyses by running three models for sperm quality parameters and four models for birds’ coloration – one for each species. In these models, we used log-transformed sperm density (three species included), logit-transformed percentage of abnormal sperm (four species included), or log-transformed intra-individual variation in sperm morphology (four species included) as dependent variables, with farming practice as the sole explanatory variable, given the limited sample size for each species.

All LMMs were run using the *lmer* function from the R package *lmerTest* (Kuznetsova et al., 2017). To avoid inflating type I error (Forstmeier and Schielzeth, 2011; Whittingham et al., 2006), we did not apply model selection, and therefore always report results for full models. Modelling assumptions (normality of residuals, normality of random effects, residuals linearity, homogeneity of variance, collinearity of factors) were checked by visual inspection of the residuals using the *check_model* function of the *performance* package (Lüdecke et al., 2021). Following recommendations in the ASA statement on p-values (Wasserstein and Lazar, 2016), we do not interpret our results based on an arbitrary threshold for statistical significance. Instead, we use p-values to measure the amount of statistical evidence against the null hypothesis and further examine effect sizes (Cohen’s *d*) along with their confidence intervals to assess the strength of the associations and the confidence we should have in those estimates.

## 3. Results

Basic descriptive statistics per species (sample sizes, median, mean ± sd, minimum and maximum) are presented in Table S1. Sample size varies based on the parameters being studied due to field constraints related to the amount of sperm that can be collected; in some cases, the quantity was too low to perform all the necessary analyses.

### 3.1. Body mass

All seven species were included in this analysis. We found no statistical evidence that bird body mass differed between organic and conventional sites (χ² = 1.33, *p* = 0.25). However, body mass was positively correlated with tarsus length and varied among species (χ² = 3.97, *p* = 0.05; χ² = 2236.70, *p* < 0.001, respectively). There was no evidence of an interaction between site and species (χ² = 4.13, *p* = 0.66). Additionally, the date of capture (Julian date) had no effect on body mass (χ² = 0.18, *p* = 0.67), but the capture session did (χ² = 8.62, *p* = 0.003). Post-hoc analyses confirmed that body mass did not differ between sites for any species (all p between 0.26 – 0.71).

### 3.2. Ornamental colorations

We included four species in the analysis (Great Tit, Cirl Bunting, Common Blackbird, Eurasian Blackcap). We found no statistical evidence that the intensity of ornamental coloration differed between farming practices or that there was an interaction between farming practice and species (χ² = 0.29, p = 0.59; χ² = 1.81, p = 0.61, respectively; Figure 1A). Likewise, there was no statistical evidence that ornamental coloration varied with body condition or Julian date (χ² = 1.00, p = 0.32; χ² = 0.00, p = 0.99). However, we did find strong statistical evidence for differences in ornamental coloration among species (χ² = 76.93, p < 0.001). Post-hoc analyses confirmed that ornamental colorations did not differ between sites for any species (all p between 0.06 – 0.89).

**Figure 1.**
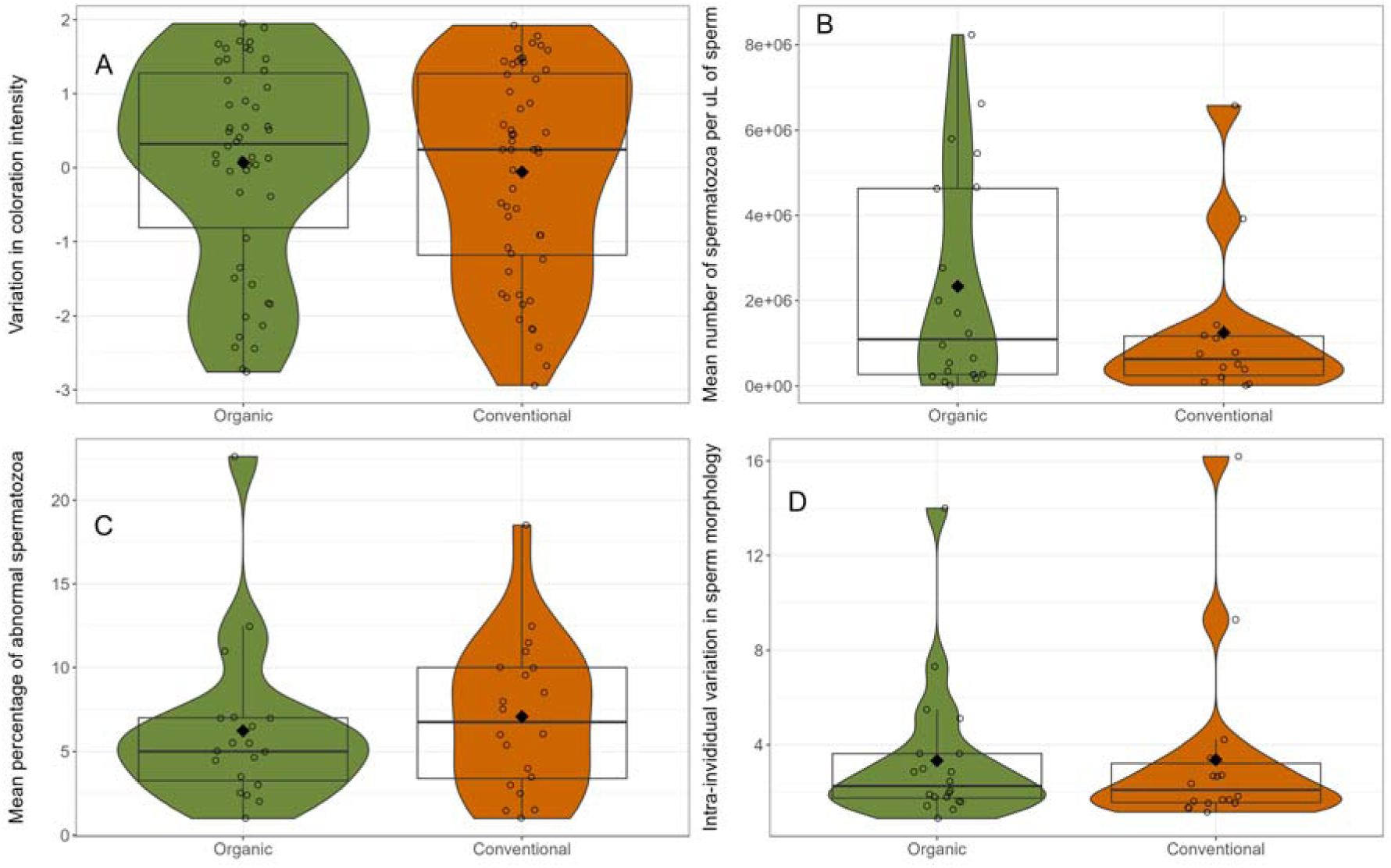
**(A)** Variation in coloration intensity in four species (Great Tit, Cirl Bunting, Common Blackbird, and Eurasian Blackcap), **(B)** sperm density in three species (Dunnock, Common Whitethroat, and Common Nightingale), **(C)** percentage of abnormal sperm in four species (Dunnock, Common Whitethroat, Common Nightingale, and Eurasian Blackcap) and **(D)** sperm morphology in four species (Dunnock, Common Whitethroat, Common Nightingale, and Eurasian Blackcap) in relation to the type of farm. Dots represent individual values. In the boxplots, means of each parameter are represented by black diamond and medians are represented by thick horizontal bars. The lower edge of the box represents the 25th percentile, while the upper edge represents the 75th percentile. Whiskers extend to 1.5 times the interquartile range. To account for inherent differences between species and the fact that each species may respond differently to the type of farming practices, the interaction between the type of farming practices and birds’ species was added in each model as a random slopes and random intercepts, respectively. It allows for each species to have a different baseline response (random intercept) and a different reaction to the change of farming practices (random slopes).

### 3.3. Sperm quality parameters

The analysis that included sperm density as the response variable was conducted on three species (Dunnock, Common Whitethroat, Common Nightingale), whereas the analyses on the percentage of abnormal spermatozoa and intra-individual variation in sperm morphology involved four species (Dunnock, Common Whitethroat, Common Nightingale, and Eurasian Blackcap). The sample size varies between parameters due to the limited volume of semen we were able to collect in the field, which prevented us from conducting all the measures of sperm quality. The results of the full models combining all species did not reveal any statistical evidence to suggest that the type of farming practice impacted any of the studied sperm quality parameters (refer to Table 1, Figure 1 B – D). Additionally, the date (Julian) had a negative influence on sperm density, but we also found small statistical evidence that sperm density decreased with birds’ body condition (Table 1). Interestingly, subgroup analyses for each species revealed medium statistical evidence for a decrease in sperm density in the Common Whitethroat and an increase in the proportion of abnormal spermatozoa in the Common Nightingale with larges effect sizes in birds living in conventional farming (p = 0.01 and p = 0.02, respectively; Figure 2). However, subgroup analyses on intra-individual variation in sperm morphology did not reveal any difference in birds living in organic and conventional farming habitats (Figure 2).

**Figure 2.**
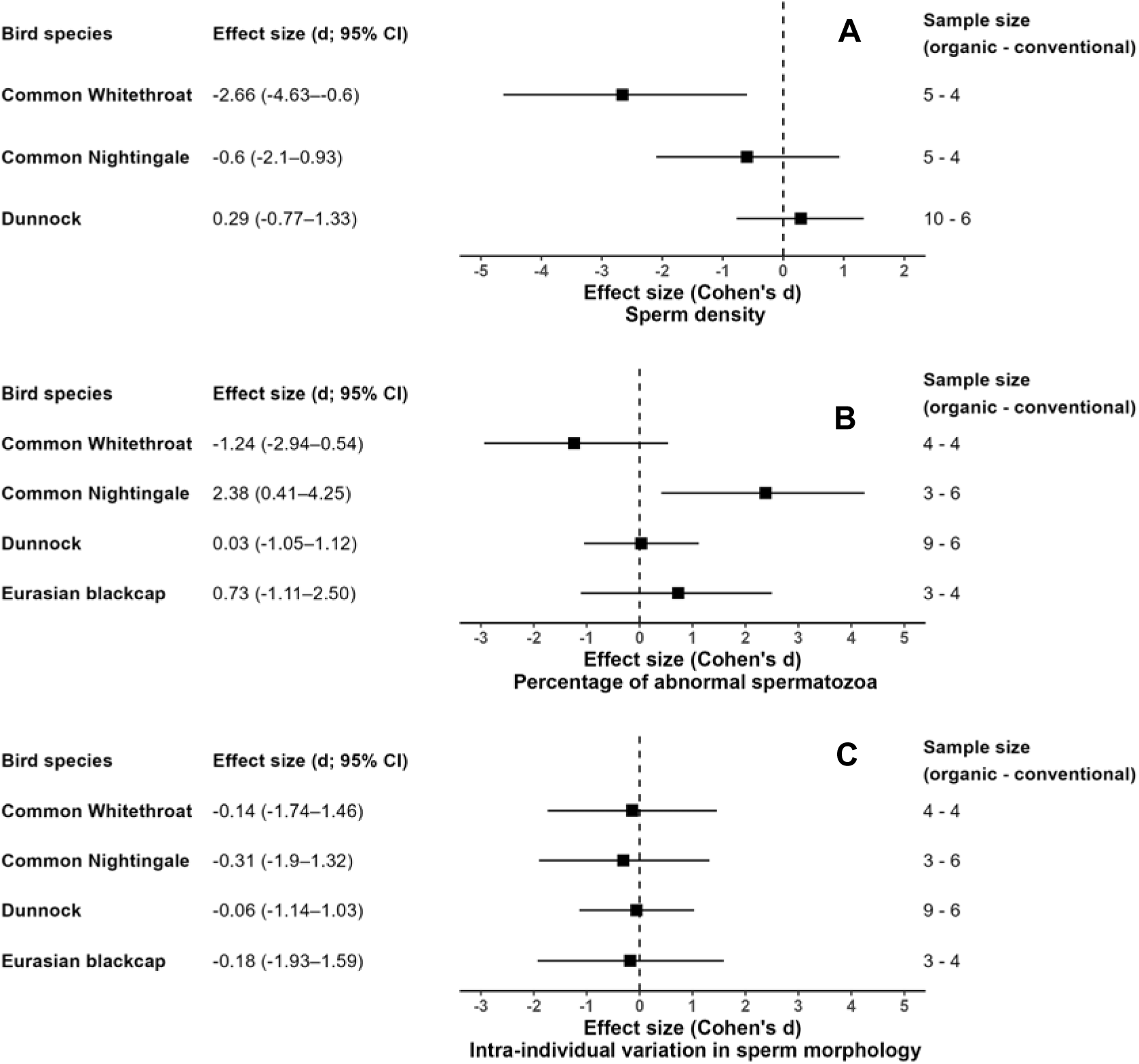
Effect sizes (Cohen’s *d*) along with their 95% confidence intervals (CI) of subgroup analyses investigating the difference in **(A)** sperm density, **(B)** percentage of abnormal spermatozoa, and **(C)** intra-individual variation in sperm morphology in the two habitats for each species. Sample sizes for each species captured in each habitat are indicated alongside the corresponding forest plots. Organic farming was set as the reference in all models. Therefore, a positive Cohen’s *d* indicates a higher value under organic farming compared to conventional farming, while a negative value indicates a lower value. A confidence interval for Cohen’s *d* that does not overlap with zero provides statistical evidence for an effect of farming practice on the parameter studied.

## 4. Discussion

In this field study, we explored the impact of conventional and organic farming environments on body mass, coloration intensity and sperm quality of several passerine birds. Contrary to our predictions, we observed no difference in either body mass or the coloration of birds caught in hedgerows within landscapes surrounded by either organically or conventionally farmed fields. Previous research conducted in the same hedgerows in 2019 has demonstrated that conventional farming negatively affects the behaviour of passerines, with birds from predominantly conventional areas showing a reduced tendency to fly away upon capture or to attack the handler (Moreau et al., 2022a). In the same study, authors did not report any difference in birds’ body condition, as calculated from the scaled mass index (Peig and Green, 2009), between conventional and organic farming. This finding, like ours, suggests that food abundance was comparable in both habitats. Moreover, both experimental and observational studies have reported the detrimental impacts of conventional food on birds’ physiology, behaviour, and reproduction, including sperm quality (Gibbons et al., 2015; Humann-Guilleminot et al., 2019; Moreau et al., 2021).

We found no evidence for a difference in sperm quality parameters between conventional and organic farming when all species were pooled. The lack of a relationship between farming practices and the sexual parameters studied (coloration and sperm quality) when combining all species might be due to several factors. First, our relatively small sample size may have reduced our statistical power to detect any effects. In addition, passerines typically have a short lifespan of 2 to 5 years in the wild and face significant reproductive costs (Bennett and Owens, 2002; Mourocq et al., 2016). Life-history trade-off hypothesis suggests that under environmental pressures, these birds might prioritize resources for reproduction over somatic maintenance, given their brief lifespan and the need to reproduce successfully (Abrams and Ludwig, 1995; Dobson and Jouventin, 2010; Kirkwood and Rose, 1991; Tazzyman et al., 2009). This prioritization could make their reproductive traits, such as sperm quality, more resilient to external stressors (e.g., pesticides) at the expense of impact on other physiological parameters.

Interestingly, subgroup analyses revealed a decrease in sperm density in the Common Whitethroat and a higher proportion of abnormal spermatozoa in the Common Nightingale, both associated with conventional farming. These two sperm quality parameters are closely linked to fertility and have been shown to be sensitive to pollutant exposure (Humann-Guilleminot et al., 2019; Humann-Guilleminot et al., 2024c; Tanga et al., 2021). A significant feature of conventional farming is the use of pesticides. Several studies conducted on birds, particularly partridges, have reported that ingestion of food containing one or more pesticides may affect both physiological parameters and reproductive success (Lopez-Antia et al., 2015a, 2015b, 2013; Moreau et al., 2021). Reduced sperm density and increased abnormalities can result from both direct and indirect disruptions to spermatogenesis. Pesticides may act as endocrine disruptors, interfering with hormone production and regulation, which in turn compromises germ cell development (Liu et al., 2025; Pandey et al., 2017). Additionally, pesticide-induced oxidative stress can directly damage spermatozoa by affecting their cellular structures and DNA, contributing to a higher proportion of abnormal sperm (Hoshi et al., 2014; Khalil et al., 2017). However, it is important to acknowledge that any environmental impacts – especially the impact of agricultural practices that involves many more aspects than pesticides ingestion – can complexly affect both survival and reproductive capacities. Therefore, it is possible that conventional farming might have a more pronounced effect on physiological aspects rather than on reproductive outcomes, which could make it difficult to detect a stronger effect in our study. In addition, while farming practices may influence various physiological traits, the specific effects on survival *versus* reproduction can vary and would need further investigation.

The lack of observable effects of habitat types on birds’ coloration and sperm quality when combining all species might also stem from the mobility of the birds studied. Although the research was conducted over a representative scale assumed to encompass a bird’s home range, we cannot exclude that birds can fly beyond the confines of the study area. Consequently, birds living in habitats surrounded by organic fields could potentially be exposed to substances from conventional farming if they fly away from their home range, or on the contrary birds could benefit from organic farming if they go beyond their intensive farming habitat. It could therefore increase the variance in our measurements, indicating a potential limitation in the study’s design. Similarly, although we included species as a random variable in each model, the increased variance in sperm quality parameters may be due to the capture of different bird species with varying mating strategies and levels of sperm competition, potentially leading to differences in sperm quality between species (Birkhead, 1998). Moreover, the differences between organic and conventional habitats may be less significant than anticipated. Research has demonstrated that contamination of untreated areas, such as organic fields, with pesticides can occur through drifting from adjacent treated fields over sometimes long distance (Botías et al., 2016; Brühl et al., 2024; Humann-Guilleminot et al., 2019; Pelosi et al., 2021). Additionally, hedgerows, although they are not directly treated, can accumulate pesticides through spray drift following the treatment of adjacent fields (Kjær et al., 2014). As a result, organic sites might still encounter some pesticide contamination, although typically at reduced levels compared to those on conventional farms. This potential contamination suggests that birds in habitats surrounded by organic farming may be not completely shielded from pesticide exposure. Conversely, it is possible that the habitats in our study were sufficiently favourable (e.g. presence of hedges, heterogeneous environment), thereby reducing significant exposure to detrimental conditions linked to agriculture. This is particularly likely since the birds were captured near hedgerows, where food sources are typically more abundant. The presence of hedgerows in agricultural landscapes is known to enhance biodiversity, potentially providing a rich food source for a variety of bird species, including those in intensive agricultural areas (García de León et al., 2021; Girard et al., 2014; Morandin et al., 2014). Consequently, any potential differences in body mass, coloration and sperm quality among the birds might have been compensated for, thus gone unnoticed. Lastly, since the effects of pesticides on breeding birds are dose-dependent (Lopez-Antia et al., 2015b; Tokumoto et al., 2013), the levels of contamination observed in our study’s sites may have been insufficient to influence the fitness components we investigated.

### Conclusion

To the best of our knowledge, this study is the first to investigate coloration intensity and sperm quality in birds captured in organic compared to conventional agricultural systems. Our results did not reveal any difference between body mass and coloration intensity between birds caught in organic *versus* conventional environments. However, despite the small sample sizes, our results indicate that conventional farming may negatively affect sperm quality through a reduction in sperm density and an increase in proportion of abnormal spermatozoa. Previous research has demonstrated the deleterious impact of intensive agriculture on various physiological, reproductive, and behavioural traits in birds. The absence of effects of the two farming systems when combining all species could stem from methodological constraints, the dilution of effects through cross-contamination of both habitats, under-exposure, or the intrinsic adaptive strategies of the species studied. Further research should aim to expand the scope to include a broader range of parameters related to both reproduction and survival, particularly including somatic physiological parameters, thus providing a more complete understanding of the impacts of farming practices on bird populations.

## Ethical statement

All experiments complied with French regulations on animal experimentation. We are grateful to the Nouvelle-Aquitaine Regional Agency of the Environment, Development and Housing for the official capture authorizations (DREAL/2019D/2323).

## Authors’ contributions

**Ségolène Humann-Guilleminot:** Methodology, Investigation, Formal analysis, Visualization, Writing - Original Draft, Writing - Review & Editing

**Audrey Bailly:** Investigation, Writing - Review & Editing

**Karine Monceau:** Writing - Review & Editing

**Mélissa Desbonnes:** Investigation, Methodology

**Vincent Bretagnolle:** Methodology, Writing - Review & Editing

**Jérôme Moreau:** Investigation, Conceptualization, Methodology, Project administration, Supervision, Writing - Review & Editing

## Acknowledgments

We thank all the farmers who granted us access to their fields. We are also grateful to everyone who assisted with bird monitoring and field data collection, and in particular to Mona Moreau for her help with field captures and data acquisition.

