## Supplementary material for "Field-based evidence of sperm quality impairment associated with conventional farming in two passerine birds"

| Table S1. Sample size, median, mean ± standard deviation (sd), minimum and maximum for body mass (“BM”; g), variation in coloration intensity (“Colo”; higher scores indicating more intense coloration), sperm density (“Sp. Dens”; mean number of spermatozoa per μL), percentage of abnormal spermatozoa (“Perc. Abn.”) and intra-individual variation in sperm morphology (“Intra-var”; μm) for each species included in each analysis. | | | | | | | | | | | | | | | |
| --- | --- | --- | --- | --- | --- | --- | --- | --- | --- | --- | --- | --- | --- | --- | --- |
|  | ***Organic*** | | | | | | | | ***Conventional*** | | | | | | |
|  | Species | *P. Modularis* | *E. Cirlus* | *C. Communis* | *S. Atricapilla* | *P. Major* | *T. Merula* | *L. Megarhynchos* | *P. Modularis* | *E. Cirlus* | *C. Communis* | *S. Atricapilla* | *P. Major* | *T. Merula* | *L. Megarhynchos* |
| BM | Sample size | 11 | 17 | 14 | 23 | 9 | 18 | 7 | 7 | 10 | 11 | 26 | 14 | 18 | 6 |
|  | Median | 18.90 | 23.60 | 13.40 | 16.30 | 18.20 | 85.15 | 20.20 | 18.60 | 24.05 | 13.50 | 16.80 | 17.80 | 85.85 | 21.05 |
|  | Mean ± sd | 18.80 ± 0.98 | 23.82 ± 1.55 | 13.36 ± 0.94 | 16.52 ± 1.67 | 18.11 ± 0.94 | 85.46 ± 5.52 | 19.89 ± 1.09 | 18.50 ± 1.32 | 23.84 ± 0.95 | 13.46 ± 0.76 | 16.68 ± 0.96 | 17.74 ± 0.52 | 86.02 ± 4.95 | 21.10 ± 1.29 |
|  | Minimum | 16.50 | 20.90 | 11.90 | 11.70 | 16.20 | 67.00 | 17.90 | 16.60 | 22.30 | 12.00 | 14.80 | 16.50 | 78.00 | 19.50 |
|  | Maximum | 19.90 | 27.30 | 15.90 | 19.90 | 19.50 | 91.60 | 20.90 | 20.80 | 25.20 | 14.60 | 19.00 | 18.30 | 95.60 | 23.10 |
| Colo | Sample size | - | 15 | - | 14 | 7 | 15 | - | - | 9 | - | 22 | 8 | 14 | - |
|  | Median | - | 0.51 | - | -1.93 | 0.29 | 1.59 | - | - | 0.50 | - | -1.56 | 0.30 | 1.48 | - |
|  | Mean ± sd | - | 0.49 ± 0.51 | - | -1.76 ± 0.91 | 0.28 ± 0.17 | 1.27 ± 0.82 | - | - | 0.65 ± 0.36 | - | -1.37 ± 0.98 | 0.33 ± 0.20 | 1.36 ± 0.68 | - |
|  | Minimum | - | -0.39 | - | -2.76 | 0.06 | -0.95 | - | - | 0.21 | - | -2.94 | -0.03 | -0.91 | - |
|  | Maximum | - | 1.44 | - | 0.30 | 0.55 | 1.95 | - | - | 1.26 | - | 1.44 | 0.58 | 1.92 | - |
| Sp. Dens. | Sample size | 10 | - | 5 | - | - | - | 5 | 6 |  | 4 |  |  |  | 4 |
|  | Median | 1.47E+06 | - | 4.63E+06 | - | - | - | 2.25E+05 | 1.31E+06 |  | 3.18E+05 |  |  |  | 2.15E+05 |
|  | Mean ± sd | 2.76E+06 ± 2.95E+06 | - | 3.52E+06 ± 2.18E+06 | - | - | - | 2.82E+05 ± 2.12E+05 | 0.02E+06 ± 0.02E+06 |  | 3.69E+03 ± 2.91E+03 |  |  |  | 3.05E+03 ± 3.59E+03 |
|  | Minimum | 1.50E+04 | - | 5.37E+05 | - | - | - | 1.00E+05 | 5.00E+05 |  | 9.00E+04 |  |  |  | 1.13E+04 |
|  | Maximum | 8.24E+06 | - | 5.80E+06 | - | - | - | 6.45E+05 | 6.58E+06 |  | 7.50E+05 |  |  |  | 7.80E+05 |
| Perc. Abn. | Sample size | 9 | - | 4 | 3 | - | - | 3 | 6 | - | 4 | 4 | - | - | 6 |
|  | Median | 6.50 | - | 8.50 | 5.00 | - | - | 2.50 | 5.50 | - | 2.75 | 9.03 | - | - | 8.00 |
|  | Mean ± sd | 5.89 ± 2.81 | - | 10.78 ± 8.85 | 4.17 ± 1.89 | - | - | 3.18 ± 1.27 | 6.00 ± 3.58 | - | 4.38 ± 4.61 | 8.01 ± 4.66 | - | - | 9.32 ± 4.95 |
|  | Minimum | 1.00 | - | 3.50 | 2.00 | - | - | 2.40 | 2.50 | - | 1.00 | 1.50 | - | - | 5.40 |
|  | Maximum | 11.00 | - | 22.60 | 5.50 | - | - | 4.65 | 11.50 | - | 11.00 | 12.50 | - | - | 18.50 |
| Intra-var | Sample size | 9 | - | 4 | 3 | - | - | 4 | 6 | - | 4 | 4 | - | - | 4 |
|  | Median | 1.90 | - | 2.92 | 1.62 | - | - | 6.40 | 1.74 | - | 2.01 | 1.44 | - | - | 6.36 |
|  | Mean ± sd | 2.13 ± 0.75 | - | 2.59 ± 1.18 | 1.72 ± 0.21 | - | - | 7.98 ± 4.13 | 2.18 ± 1.12 | - | 2.25 ± 0.83 | 1.72 ± 0.64 | - | - | 7.91 ± 6.26 |
|  | Minimum | 1.27 | - | 0.89 | 1.57 | - | - | 5.11 | 1.16 | - | 1.60 | 1.32 | - | - | 2.71 |
|  | Maximum | 3.63 | - | 3.63 | 1.96 | - | - | 14.01 | 4.21 | - | 3.38 | 2.67 | - | - | 16.19 |
